# CD13 Activation Assembles Signaling Complexes that Promote the Formation of Tunneling Nanotubes in Endothelial Cells

**DOI:** 10.1101/2024.05.10.593402

**Authors:** Emily Meredith, Brian Aguilera, Fraser McGurk, Pengyu Zong, Lixia Yue, Mallika Ghosh, Linda H Shapiro

**Author notes:** Corresponding authors: Mallika Ghosh, PhD Linda H Shapiro, PhD, Center for Vascular Biology, University of Connecticut School of Medicine Farmington, CT 06030.

## Abstract

Transmembrane CD13 assembles protein complexes at the plasma membrane to enable diverse cellular processes such as cell-cell adhesion, focal adhesion turnover, endocytosis and recycling of cell surface proteins. In this study, we demonstrate a novel CD13-dependent assembly platform that regulates phosphoinositide (PI) signal transduction during the formation of Tunneling Nanotubes (TNTs). TNTs are actin-based, membrane-delimited bridges that facilitate intercellular communication by connecting distant cells to physically transfer subcellular cargoes. TNTs form between various cell types under stress conditions, but few molecular TNT-inducers exist. Human Kaposi’s sarcoma-derived endothelial cells (KSECs) readily form stress-induced TNTs capable of transferring Ca^2+^ and membrane molecules between cells, with clear accumulation of CD13 and actin at the base of the protrusions. Alternatively, CD13-null KSECs form fewer TNTs and Ca^2+^ transfer is markedly reduced. Mechanistically, CD13-mediated TNT formation requires activation of CD13, Src, FAK and Cdc42 to allow tethering of the IQGAP1 and ARF6 complex at the membrane to activate the phosphatidylinositol-4-phosphate-5-kinase PI5K. This increases local phosphatidylinositol 4,5-bisphosphate (PI(4,5)P2) levels to promote the actin-polymerization and membrane protrusion necessary for TNT formation. Therefore, CD13 is a novel molecular PIP regulator and TNT trigger that will facilitate the dissection of downstream pathways and mechanisms regulating TNT formation.

## Introduction

Acquiring and exchanging molecular information is essential for the survival of multicellular organisms. Communication between cells as well as cells with their microenvironment is fundamental to preserving tissue homeostasis and restoring normal function upon damage or deterioration. Logically, a high degree of specificity is required for accurate transfer; it is vital that a particular initiating cell communicate the appropriate message to the correct recipient (Yamashita *et al*., 2018). While numerous processes have been described that mediate cell-cell communication such as the diffusion of secreted soluble molecules or the release of microvesicles or cargo-loaded exosomes, these lack the strict specificity or reliability required in some cases (Wolpert, 2016; Caviglia and Ober, 2018). Indeed, the microenvironment of damaged tissues is highly toxic and proteolytic, yet transplanted mesenchymal stem cells are able to transfer healthy organelles and vesicles to rescue damaged cells, suggesting a more ‘protected’ means of communication must exist. Alternatively, protrusion-based cellular interactions such as filopodia (Gardel *et al*., 2010; Jacquemet *et al*., 2015), cytonemes (Kornberg and Roy, 2014; Stanganello and Scholpp, 2016) and although less well studied, the tunneling nanotubes (TNTs) (Onfelt *et al*., 2004; Rustom *et al*., 2004; Baker, 2017; Yamashita *et al*., 2018; Hanna *et al*., 2019; Genna *et al*., 2023) would provide specificity over soluble- or exosome-based intercellular communication mechanisms (Caviglia and Ober, 2018; Yamashita *et al*., 2018). Filopodia are short, thin, actin-containing protrusions localized at the cell base with important roles in sensory and cell-cell communication (Jacinto and Wolpert, 2001; Fischer *et al*., 2019), while cytonemes are long, close-ended specialized filopodia that connect distant cells to transport signals between various cell types in Drosophila and zebrafish (Gonzalez-Mendez *et al*., 2019), (Eom *et al*., 2015), (Hamada *et al*., 2014), (Mattes *et al*., 2018). Similar to cytonemes, TNTs are long, thin, actin-based membrane protrusions that are open-ended and connect distant cells to mediate the tightly regulated, targeted transfer of signals, endosomes, organelles and proteins between homotypic or heterotypic cells (Yasuda *et al*., 2011; Liu *et al*., 2014; Wolpert, 2016; Vignais *et al*., 2017; Caviglia and Ober, 2018; Yamashita *et al*., 2018; Hanna *et al*., 2019; Liu *et al*., 2019; Mittal *et al*., 2019b) in a number of cell and tissue contexts (Gerhardt and Betsholtz, 2003; Pober and Sessa, 2007; Wittchen, 2009; Vestweber *et al*., 2013; Reglero-Real *et al*., 2016; Caporali *et al*., 2017; Johnson *et al*., 2017). These combined observations support a mechanism whereby protrusion-based structures enable highly specific, long-range intercellular communication and may explain the propagation of signals through complex and potentially lethal environments to influence cell fate *in vivo*.

Despite their initial description nearly 20 years ago (Rustom *et al*., 2004), the extracellular and intracellular signals and specific molecules that either trigger or specifically inhibit the induction, formation and function of these protrusions are largely unknown (Belian *et al*., 2023). TNTs appear to form in response to metabolic stress such as deprivation of serum proteins and high glucose conditions, but few specific molecules have been identified that induce TNT formation (Yamashita *et al*., 2018; Belian *et al*., 2023). Logically, both the membrane and the cytoskeleton must play a role in protrusion formation. Indeed, inhibition of actin polymerization abrogates TNT formation and various actin-binding, membrane tethering, signal regulators and actin-regulatory proteins have been implicated in TNT formation and assembly such as Cdc42, IQGAP1, LST1, RalA, M-sec and filamin in various cell types (Hase *et al*., 2009; Schiller *et al*., 2013; Delage *et al*., 2016; Grabowska *et al*., 2020; Barutta *et al*., 2021; D’Aloia *et al*., 2021). Similarly, membrane organization and function are controlled by membrane lipids that sort the membrane into dynamic subdomains, assembling signaling platforms (Takenawa and Itoh, 2001; Wenk, 2005; Simons and Sampaio, 2011; Delage and Zurzolo, 2013). Membrane lipids themselves, particularly phosphatidylinositol or sphingolipid species, also induce signaling cascades and regulate activity or membrane localization of various proteins via direct interaction to impact many cellular functions (Takenawa and Itoh, 2001; Chichili and Rodgers, 2009; Saarikangas *et al*., 2010; Delage and Zurzolo, 2013; Regen, 2020). Emerging evidence indicates that the cytoskeletal makeup and appearance of TNTs is highly-context and cell-type dependent which, combined with the lack of specific markers, makes it a challenge to definitively distinguish among the various types of protrusions (Naphade *et al*., 2015; Delage *et al*., 2016; Caviglia and Ober, 2018; Yamashita *et al*., 2018; Belian *et al*., 2023). Similarly, the mechanisms guiding TNT formation show a fair degree of cell specificity and whether a common, unifying process exists remains unclear. Further investigation is clearly warranted to provide tools to answer these questions and to fully exploit these novel mechanisms of cell communication.

Our observations have established that the transmembrane molecule CD13 is remarkably multifunctional and regulates diverse processes such as filopodia formation, mesenchymal stem cell (MSC) rescue, maintenance of the stem cell niche, endocytic trafficking, angiogenesis, membrane organization, cell-cell fusion and cell-matrix and cell-cell adhesion (Pasqualini *et al*., 2000; Bhagwat *et al*., 2001; Mina-Osorio *et al*., 2006; Petrovic *et al*., 2007; Pereira *et al*., 2013; Subramani *et al*., 2013; Ghosh *et al*., 2014; Rahman *et al*., 2014a; Rahman *et al*., 2014b; Ghosh *et al*., 2015; Ghosh *et al*., 2019; Ghosh *et al*., 2021). Indeed, our recent studies demonstrated that CD13 recruits a complex containing IQGAP1/ARF6/EFA6 to the plasma membrane and tethers this complex to the actin cytoskeleton via α-actinin (Ghosh *et al*., 2019). In the present study, we show that CD13 is required for the formation of long, thin protrusions above the substrate that connect distant cells and contain actin and important signaling molecules, consistent with TNTs (Chakraborty *et al*., 2023). We further illustrate that CD13 expression is necessary for the transfer of calcium signals and membrane molecules, which occurs in WT but not in CD13^KO^ KSECs. Interestingly, activation of CD13 by the crosslinking mAb 452 enhances TNT formation and cells engineered to lack CD13 form significantly fewer TNTs, suggesting an active role for CD13 in TNT formation. Morphologically, CD13 and actin are both highly abundant and co-localize at the base as well as along the length of the TNTs under both stressed and induced conditions, consistent with CD13 contributing to actin organization in TNTs. CD13-interacting proteins such as α-actinin, and IQGAP1 are mislocalized in the absence of CD13 with accumulation of ARF6, IQGAP1, and specific PIP species PI(4,5)P at the base and/or along the length of TNTs in WT KSECs. To determine the potential CD13 effects on membrane lipid levels, we demonstrated that treatment with CD13-crosslinking mAb 452 increased PIP5K activity and PI(4,5)P2 levels in WT but not in CD13^KO^ cells, thus establishing a novel connection between CD13 and lipid signaling. Mechanistically, treatment with peptides designed to interfere with CD13’s cytoplasmic signaling domain diminished TNT formation, as did inhibition of IQGAP1, Cdc42, FAK, or Src signaling (Subramani *et al*., 2013), suggesting that CD13 activation-dependent signal transduction mediates TNT formation. Collectively, these investigations support the notion that CD13 is a unique regulatory node that coordinates the assembly, location and function of intracellular signaling complexes to promote various forms of intercellular communication.

## Results

### CD13 expression facilitates TNT formation

We previously showed that CD13 regulates endothelial cell invasion by triggering Cdc42-dependent formation of long actin-positive protrusions that were similar in morphology to structures named TNTs (Chakraborty *et al*., 2023). Furthermore, our recent studies indicated that CD13 can act as a linker between the plasma membrane and the actin cytoskeleton (Ghosh *et al*., 2019), suggesting a potential connection between CD13 and TNT formation in endothelial cells. We induced TNT formation in the human Kaposi’s sarcoma-derived endothelial cell line (KSEC) by serum deprivation and observed abundant CD13 expression along the length of actin-containing TNTs (white arrows, Figure 1A; Figure S1), consistent with a role for CD13 in TNT formation and/or function. To further investigate this notion, we CRISPR-engineered KSEC cells to delete CD13 (CD13^KO^ KSECs) or scrambled guide RNA controls (WT KSECs) and cloned and tested independent lines (Ghosh *et al*., 2019). The number of potential TNTs/cell (TNT index) (Desir *et al*., 2018; Dubois *et al*., 2018) in WT KSECs increased under metabolic stress conditions induced by serum starvation (SS) or high glucose (HG) for 1h (4.5g/L glucose, CM vs. SS/HG; 3.5 vs. 2.5, Figure 1B) as compared with complete medium control, while the TNT index of CD13^KO^ KSECs was significantly lower under all conditions tested, confirming that CD13 contributes to what appears to be TNT formation in endothelial cells. Few specific markers of TNT currently exist, and thus their identification relies on their characteristic location above the substrate and their ability to transfer cargo from one cell to another (Chakraborty *et al*., 2023). Examination of z-stack images indicated that the majority of protrusions from CD13^KO^ KSEC were restricted to an interval of less than a micron above the substrate, while WT protrusions spanned distances up to 2.88 microns above the culture surface (starting at 0.24μm, Z-stacks were taken in 0.24 μm intervals for a total of 2.64 μm in total stack height, with each 0.24 μm section considered a bin). Protrusions spanning multiple z-sections were assigned to the bin containing the majority of its length. Long, actin-containing protrusions present in the middle sections of the z-stack (1.3 μm -2.4 μm above the substrate) were identified as TNTs (Figure 1C, S2). These were also appreciably longer than in CD13^KO^ cells (15 μm vs. 10 μm) (Figure 1D, consistent with TNT-like protrusions (Chakraborty *et al*., 2023). Since TNTs can be either actin or microtubule-based (Resnik *et al*., 2022), we treated WT and CD13^KO^ KSECs with microtubule (nocodazole) or actin (Cytochalasin B, D) polymerization inhibitors or the Cdc42 GTPase inhibitor (ML141) and measured the TNT index. We found that treatment with actin and Cdc42, but not microtubule inhibitors significantly decreased TNT formation in WT KSECs (Figure 1E, F), indicating that the KSEC protrusions are largely actin-driven, consistent with actin-based endothelial TNTs (Hashimoto *et al*., 2016; Dupont *et al*., 2018; Belian *et al*., 2023).

**FIGURE 1.**
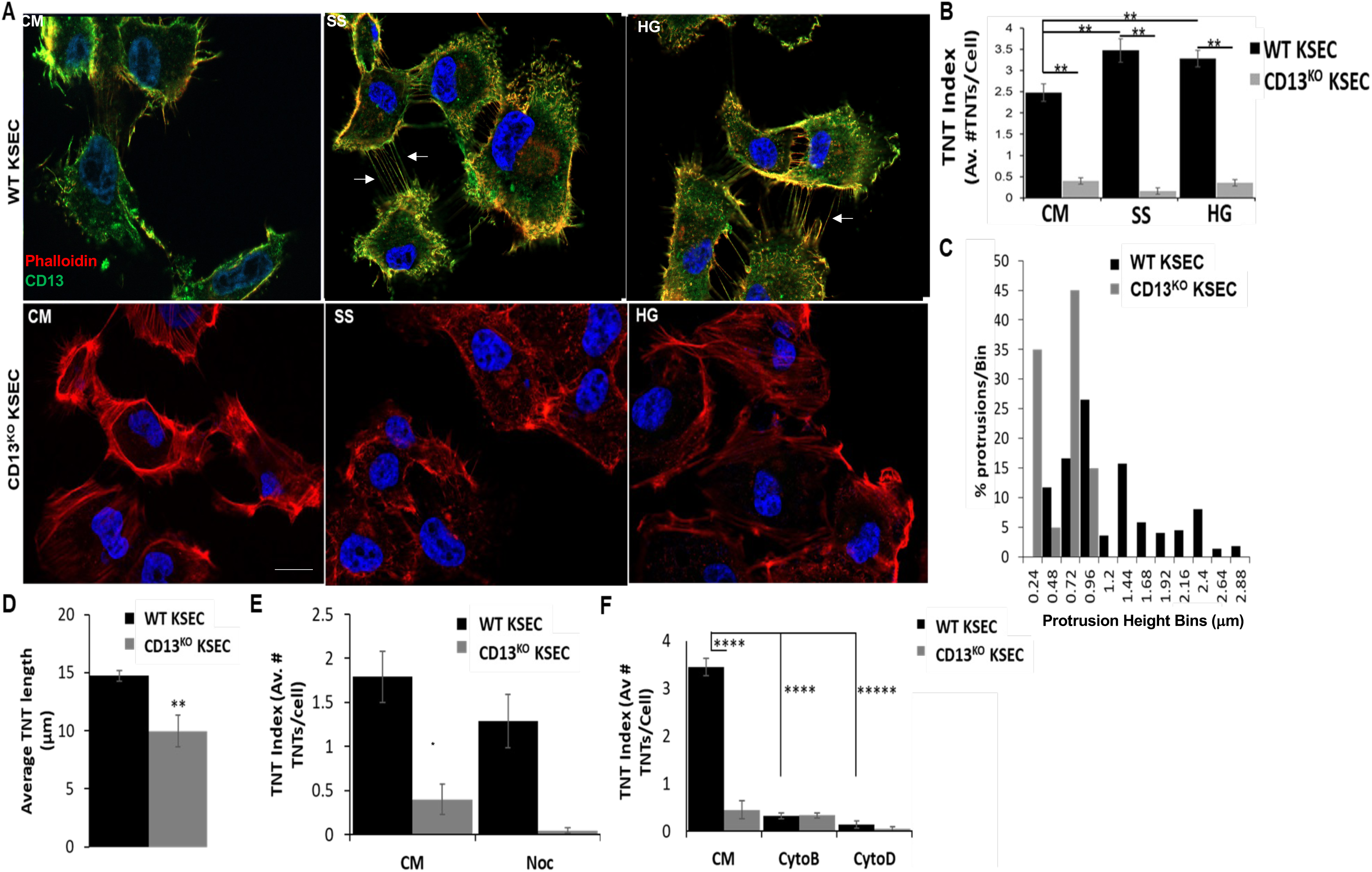
CD13 is required for TNT formation under metabolic stress. **(A)** WT and CD13^KO^ KS1767 (KSEC) cells were incubated under stress conditions (SS; serum starvation, HG; high glucose) for 1hr. WT KSEC TNTs stained positive for actin (phalloidin; red), and CD13 (green). White arrows highlight TNTs. Scale bar: 10μm. **(B)** The TNT index was calculated by dividing the total number of TNTs within a FOV by the total number of cells. WT cells form significantly more TNTs under stress conditions compared to CD13^KO^ KSECs when compared to complete media (CM) control. **(C)** the elevation of TNTs from the substrate in WT and CD13^KO^ cells was quantified in a series of 0.24μm height bins represented as percent TNTs per bin. Phalloidin^+^ protrusions present in the middle section of the z-stacks were identified as TNTs. **(E)** Treatment of WT and CD13^KO^ cells with nocodazole, a microtubule inhibitor (150nM) for 4hrs prior to fixation and TNT quantification, supported by **(F)** a significant decrease in TNT formation following treatment with actin inhibitors CytoD (250nM), CytoB (250nM). Data are represented as mean ± standard error. *p≤0.05.

### CD13-dependent protrusions are functional TNTs

To examine whether CD13-dependent TNTs have the ability to form functional connections that transfer cargo between cells (Watkins and Salter, 2005; Wang *et al*., 2010; Smith *et al*., 2011; Wang and Gerdes, 2015; Lock *et al*., 2016; Alarcon-Martinez *et al*., 2020), we performed Ca^2+^ transfer experiments using real-time Fura2-AM Ca^2+^ imaging. Mechanical stimulation of cells elicits robust cytosolic Ca^2+^ elevations that can be propagated to physically separated cells via TNTs (Belian *et al*., 2023). In our study, WT and CD13KO KSECs were loaded with Fura-2 AM and mechanical stimulation with a patch electrode induced an increase in the intracellular Ca^2+^ concentration in both WT and CD13^KO^ cells (Figure 2A, red cell ‘touching’). Positive transfer of the intracellular Ca^2+^ flux to surrounding cells occurred at a significantly faster rate in the WT compared to CD13^KO^ KSEC group, showing a significantly higher [Ca^2+^] that was sustained for 120 sec post perturbation, eventually equalizing by the 600 sec time point (Figure 2A-D). Enlargement of the images illustrates that the Ca^2+^ transfer was facilitated by TNTs in WT but not in CD13^KO^ KSECs (Figure 2E). This data indicates that the CD13-dependent protrusions are functional and capable of transferring Ca^2+^, and that the efficiency of Ca^2+^ transfer is compromised in the absence of CD13, consistent with TNTs

**FIGURE 2.**
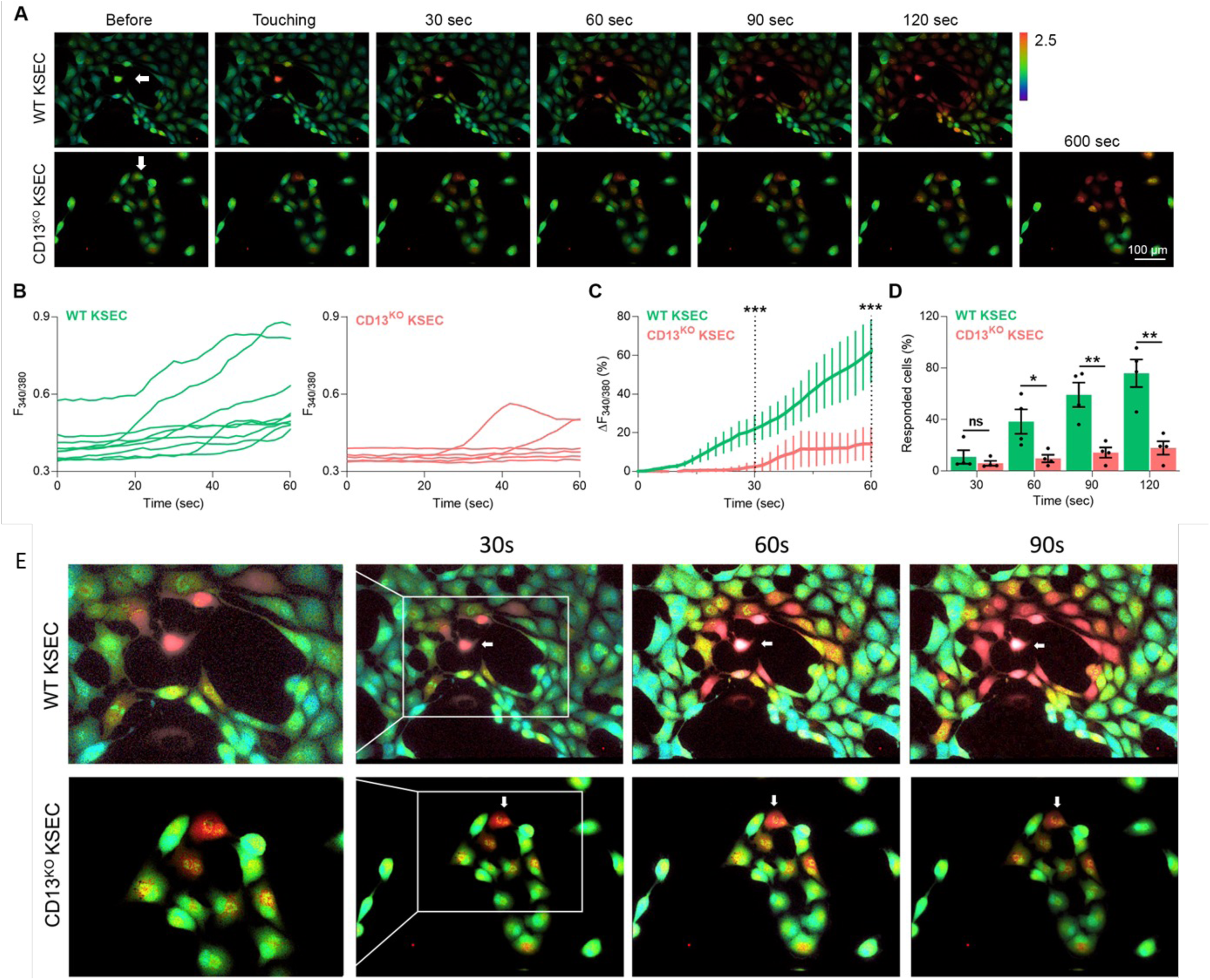
CD13-dependent TNTs functionally transfer Ca^2+^. **(A)** WT and CD13^KO^ KSECs were loaded with Fura-2 AM (2.5 µM) and mechanically stimulated with a patch electrode to monitor Ca^2+^ transfer between cells over 120 sec. White arrows denote the stimulated cell. **(B)** Individual cell traces of Ca^2+^ flux transferred to neighboring cells in cultures of WT or CD13^KO^ KSECs following stimulation. **(C)** Averaged cell traces of WT and CD13^KO^ KSECs showed a significant decrease in Ca^2+^ flux in CD13^KO^ KSECs. **(D)** Quantification of the number of responsive cells (cells with increased Ca^2+^ content) in WT and CD13^KO^ KSEC cultures. **(E)** Enlarged representative images of WT and CD13^KO^ KSEC cultures (from **2A)** demonstrate that Ca^2+^ transfer between cells is propagated via TNTs. Data are represented as mean ± standard error. *p<0.05. Scale bar is 100μm.

### CD13 activation specifically induces TNT formation

Cell surface receptor activation is often triggered by treatment with specific crosslinking monoclonal antibodies (Florey *et al*., 2010; Liu *et al*., 2012). We have shown that in monocytic cells CD13 is activated by the crosslinking mAb 452, resulting in a rapid, Src-dependent phosphorylation of tyrosine 6 (Y^6^) on CD13’s cytoplasmic tail (Subramani *et al*., 2013). CD13 activation coordinates its subsequent interaction with signaling mediators and actin-binding proteins, ultimately connecting the actin cytoskeleton to the plasma membrane to promote cell-cell and cell-extracellular matrix adhesion (Mina-Osorio *et al*., 2008; Subramani *et al*., 2013). To investigate the effect of CD13 cross-linking/activation on TNT formation, we treated WT or CD13^KO^ KSECs with the mAb 452 (1.8μg/mL, 1hr) and found a significant induction in TNT formation as compared to vehicle or serum-stressed controls, suggesting that not only is CD13 necessary for TNT formation, but its activation markedly promotes it (Figure 3A-B and S3). By contrast, treatment with either a non-activating anti-CD13 antibody (WM-47, 0.1μg/mL) (Mina-Osorio *et al*., 2006), or an antibody that binds and activates a distinct endothelial cell-surface protein CD31 (Berman and Muller, 1995) (Figure 3, CD31-crosslink, 1μg/mL) showed minimal effect on WT KSEC TNT formation, emphasizing that specific activation of CD13, but not cell surface molecules in general, underlies TNT formation. Inhibition of CD13’s enzymatic activity by treatment of KSECs with Bestatin (100μg/mL) also showed no significant effect on TNT formation, indicating that CD13-mediated TNT formation is independent of its aminopeptidase activity (Ghosh *et al*., 2012; Ghosh *et al*., 2015) (Figure 3B).

**FIGURE 3.**
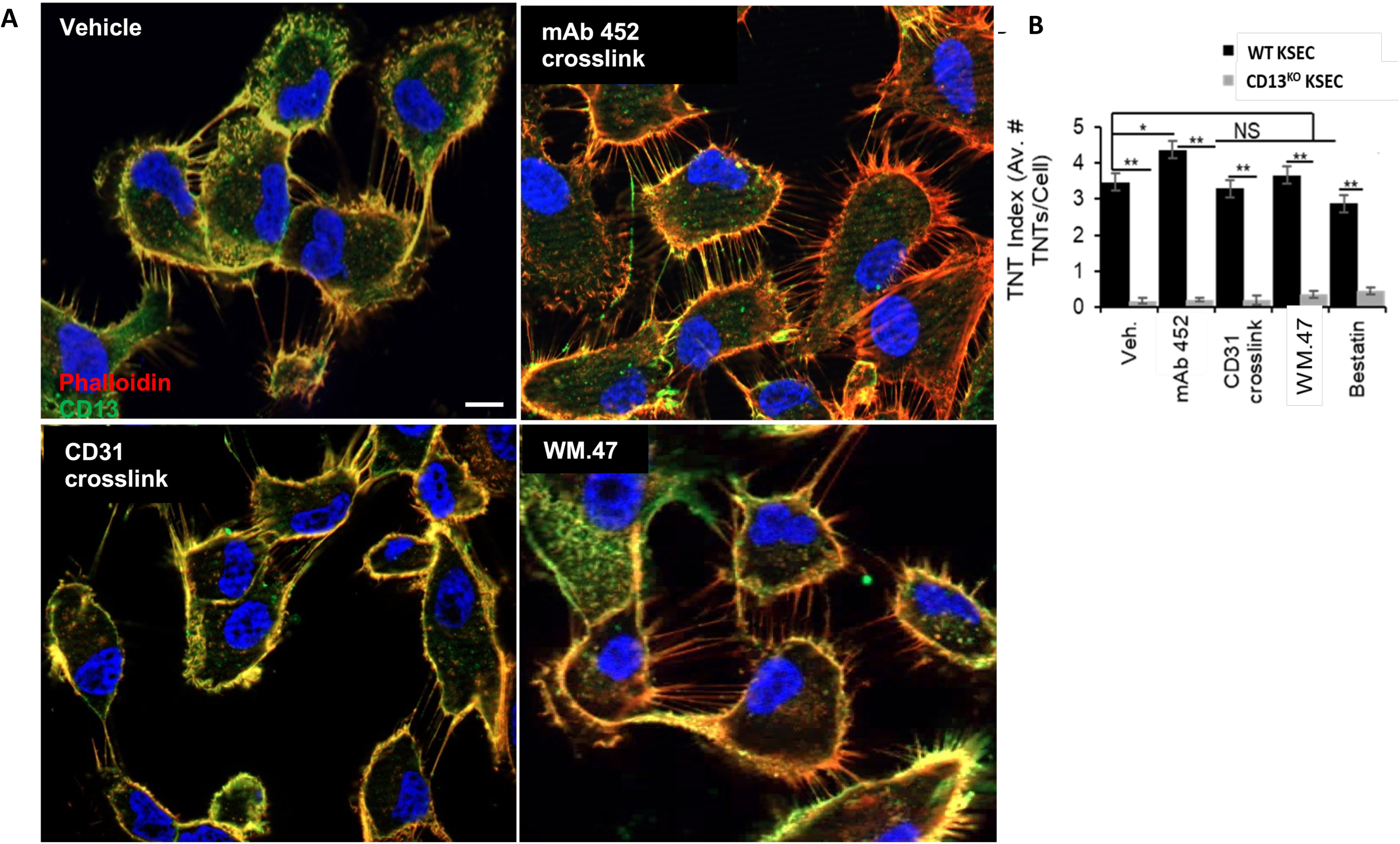
Induction of TNT formation is specific to activation of CD13. **(A)** WT KSECs were treated with indicated mAbs or control for 1hr prior to staining for actin (phalloidin; red), CD13 (green) and DAPI (blue). Arrows highlight TNTs. **(B)** Quantification of TNTs in WT and CD13^KO^ KSECs in all treatment conditions: vehicle control, CD13-activating mAb 452 (1.8μg/mL), CD31-activating mAb (1.0μg/mL), CD13 non-activating mAb WM-47 (0.1μg/mL) or bestatin (100μg/mL; an inhibitor of CD13’s protease activity). Scale bar: 10μm. Data are represented as mean ± standard error. *p<0.05.

### CD13 co-localizes with actin at the base of TNTs

A marked colocalization of CD13 and actin expression was evident at the base of TNT protrusions on the membrane. To compare the overall intensity of expression of CD13 and actin at various points along the membrane, we performed line scans by measuring the pixel intensities of either anti-CD13 or phalloidin stains in increments along the entire cell membrane by ImageJ software. (Figure 4A-B). Representative overlays normalized to the peak average pixel intensity of CD13 (black) and phalloidin (blue) line scans are shown in Figure 4C-D. The relative distance between lines on the graphs corresponds to the differences in intensity between CD13 and phalloidin stains in the same area of the membrane. These measurements indicated that intensities of both CD13 and actin were significantly higher at the base of the TNTs than at membrane areas without TNTs in serum starved conditions, which was further enhanced by mAb 452 treatment (Figure 4C-E). Furthermore, CD13 expression also correlated with cholesterol (filipin stain) in SS conditions (r = 0.68), which again significantly increased upon mAb 452 treatment (r = 0.85) and the curves noticeably aligned more closely, likely due to CD13 activation-mediated cytoskeletal rearrangement (Figure S4A-E). Similarly, staining with the lipid raft marker Cholera Toxin B (CTB) was significantly higher in WT membrane at the site of TNT initiation and along the length of the TNT than CD13^KO^ (Figure S4F-G). Taken together, this data suggests that CD13 activation coordinates the assembly of actin and cholesterol to promote TNT formation, prompting further investigation into potential mechanisms underlying CD13-mediated TNT formation.

**FIGURE 4.**
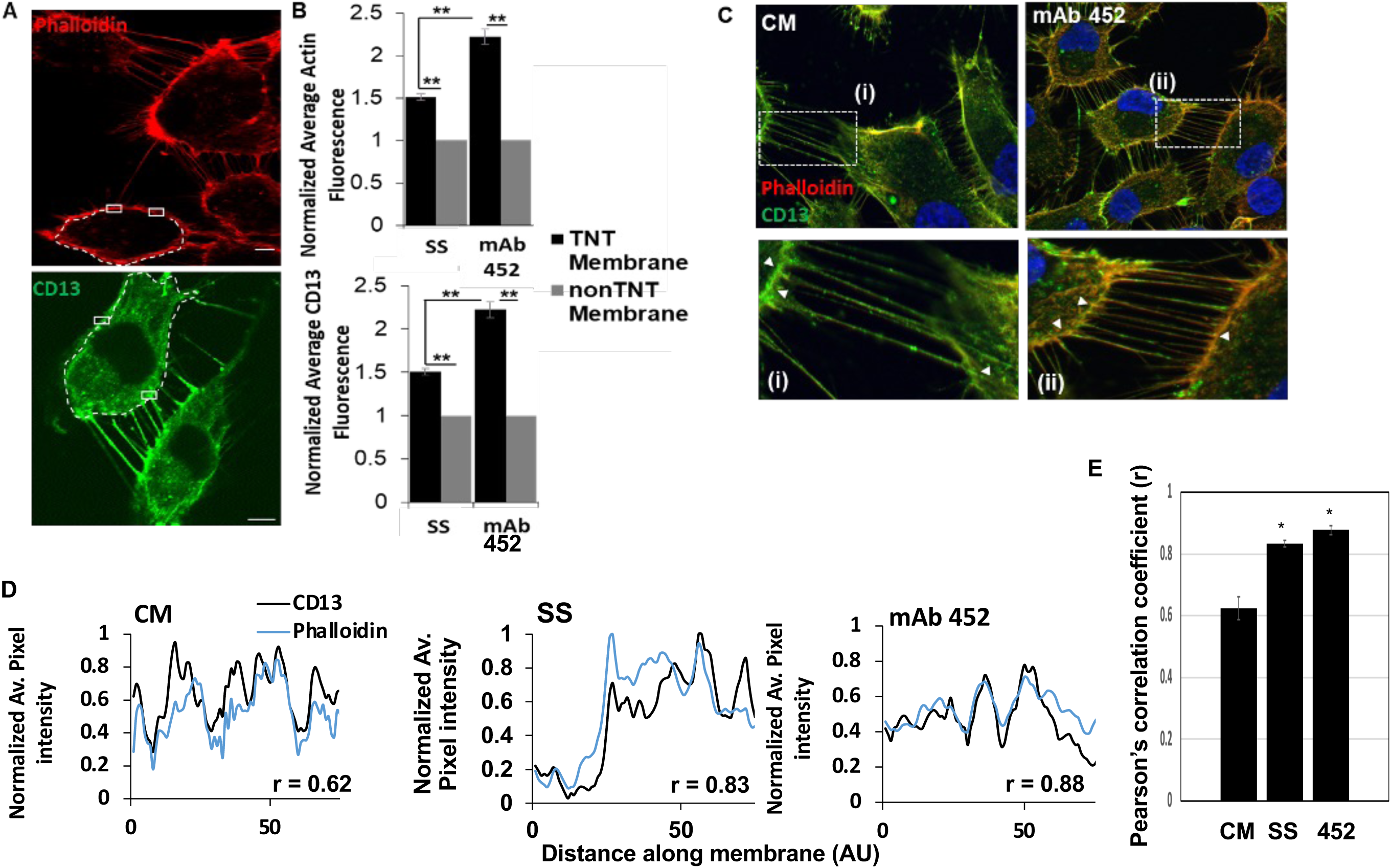
CD13 and actin co-localize at the base of TNTs. **(A)** Representative images of actin (red; top) or CD13 (green; bottom) staining of WT KSEC cells. **(B)** Quantification of the normalized average fluorescence measurements for both actin and CD13, calculated using cells entirely contained within a single FOV. Membranes were delimited by systematically measuring the fluorescence in uniformly sized areas (white boxes) around the entire membrane. Areas containing TNTs were classified as ‘TNT Membrane’, while the others, ‘non-TNT membrane’. **(C)** Representative images and **(D)** line scans of normalized average pixel intensity of CD13 (black) and phalloidin (blue) under complete media (r = 0.62), serum-starved (r = 0.83) or mAb 452 (r = 0.88) treatment conditions. **(E)** Colocalization was calculated using the Pearson Correlation Coefficient (r) for CD13 and phalloidin under complete media (cm) or 452 mAb treatment conditions. Scale bar: 10μm. Data are represented as mean ± standard error. *p<0.05.

### CD13’s cytoplasmic tail coordinates TNT formation

We have shown that although CD13’s cytoplasmic tail contains only 8 amino acids, it can interact with a number of other membrane-tethered and cytoplasmic proteins (Subramani *et al*., 2013) and blocking these interactions with interfering peptides impairs subsequent CD13-dependent functions, such as cell migration (Subramani *et al*., 2013; Ghosh *et al*., 2019; Ghosh *et al*., 2021). To determine whether TNT formation involves CD13’s cytoplasmic tail, we loaded KSECs with the CD13-blocking (MAKGFYI; 1μg/mL) or a control peptide (KFMYGAI; 1μg/mL) using BioPORTER. Blocking the CD13 cytoplasmic tail interactions resulted in a reduction in TNTs as compared to control peptide or vehicle (Figure 5A-B: Figure S5A) but did not affect CD13 expression levels (Figure 5C). The migration of cells transfected with the CD13-blocking peptide into the open area was also significantly impaired compared to control peptide in an *in vitro* scratch assay (Figure S5B), confirming that the peptide blockade is functioning correctly. Therefore, CD13-dependent TNT formation is mediated via protein interaction with its cytoplasmic tail.

**FIGURE 5.**
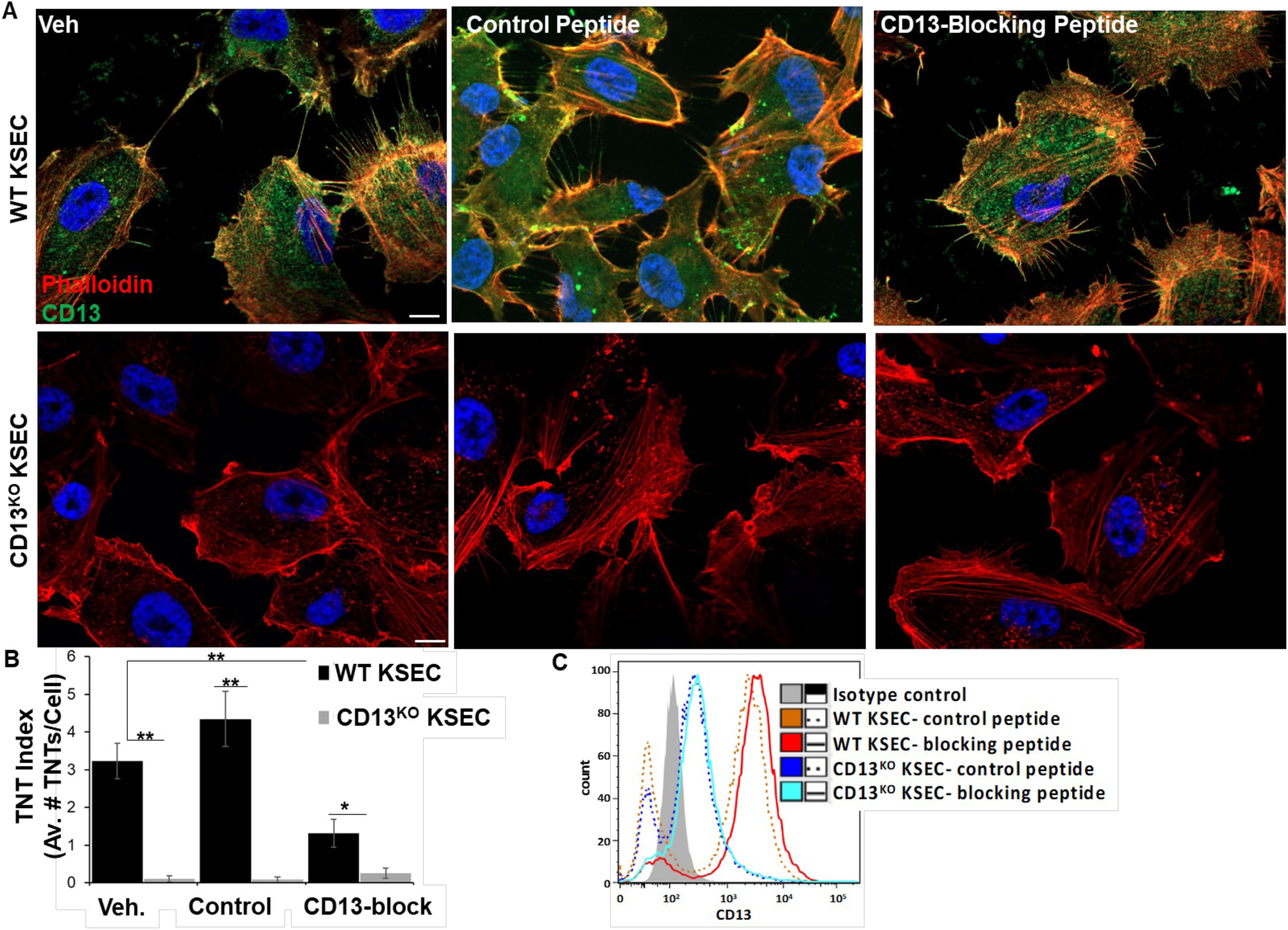
CD13’s cytoplasmic tail is required for TNT formation. **(A)** WT and CD13^KO^ KSECs were loaded with the CD13-blocking or control (1μg/mL) peptide and stained for actin (phalloidin; red), CD13 (green) and DAPI (blue) and TNT formation quantified in **(B)**. **(C)** Flow cytometric analysis of CD13 surface expression on peptide-loaded WT and CD13^KO^ KSECs shows neither peptide had a significant impact on CD13 surface expression. Figure S5 confirms that the CD13 blocking peptide is functioning correctly. Scale bar: 10μm. Data are represented as mean ± standard error. *p<0.05.

### CD13-mediated TNT Formation Involves FAK/Src Signaling, small GTPases and PI(4,5)P2 Generation

We have demonstrated that various protein interactions with CD13’s cytoplasmic tail can occur in both constitutive and activation-dependent manners, triggering overlapping and complex signaling cascades to control diverse cell functions (Mina-Osorio *et al*., 2008; Subramani *et al*., 2013; Ghosh *et al*., 2014; Rahman *et al*., 2014a; Rahman *et al*., 2014b; Ghosh *et al*., 2019; Ghosh *et al*., 2021). Intriguingly, two of these pathways have been previously implicated in TNT formation by other groups: FAK/Src signaling (Saenz-de-Santa-Maria *et al*., 2017; Tishchenko *et al*., 2020) and activation of the ARF6 small GTPase (Bhat *et al*., 2020). We had shown that the phosphorylation of FAK and Src kinases is reduced in CD13^KO^ compared to resting WT cells and in CD13^KO^ mice *in vivo* in a skeletal muscle ischemic injury model (Subramani *et al*., 2013; Rahman *et al*., 2014a; Rahman *et al*., 2014b) and that activation of CD13 in human monocytic cell lines induces marked stimulation of these kinases, prominent alterations in the actin cytoskeleton and increased cell-cell and cell-ECM adhesion (Mina-Osorio *et al*., 2008; Subramani *et al*., 2013; Ghosh *et al*., 2014; Rahman *et al*., 2014a; Rahman *et al*., 2014b; Ghosh *et al*., 2019; Ghosh *et al*., 2021). Treatment of KSEC with inhibitors of FAK Y397 (FAK14; 20μM) and Src Y416 (PP2; 10μM) phosphorylation for 1h abrogated TNT formation in WT to the level of the CD13^KO^ KSECs (Figure 6D, S6). This phenotype could not be rescued by treatment with the activating mAb 452, implying that CD13 mediates TNT formation via FAK-Src signal transduction pathways downstream of CD13 activation (Figure 6D).

**FIGURE 6.**
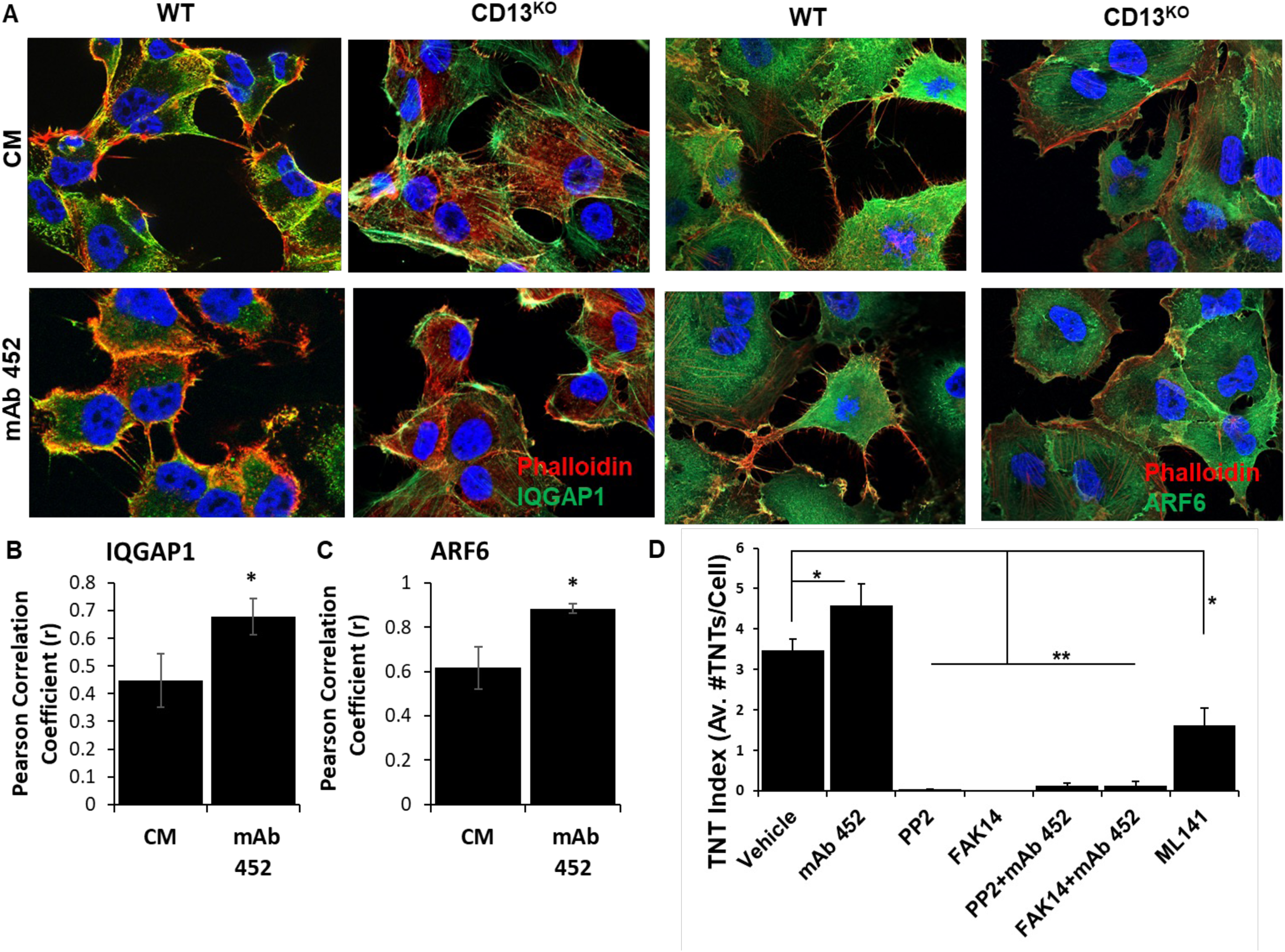
IQGAP1 and ARF6 are mislocalized in the absence of CD13. **(A)** Representative images showing IQGAP1 (green, left two panel) or ARF6 (green, right two panel) and phalloidin (actin; red) staining under control (CM) or activating (mAb 452; 1.8μg/mL) conditions. Quantification of colocalization between **(B)** IQGAP1 and phalloidin or **(C)** ARF6 and phalloidin upon CM or mAb 452 treatment as calculated by Pearson’s coefficient. **(D)** Treatment with inhibitors of major actin-organizing proteins such as FAK (FAK14; 20μM), Src (PP2; 10μM) or CDC42 (ML141; 20μM). Data are represented as mean ± standard error. *p<0.05.

CD13 also directly regulates endocytosis and endosomal trafficking by tethering a complex containing the scaffolding protein IQGAP1, the ARF6 member of the Arf family of GTPases and the actin-binding protein α-actinin to the plasma membrane (Ghosh *et al*., 2019). In the absence of CD13, these proteins are mislocalized, resulting in aberrant recycling of cell surface proteins (Ghosh *et al*., 2019). Independent studies (Bhat *et al*., 2020) demonstrated that in a neuronal cell line, ARF6 participates in a Rab GTPase signaling cascade that results in TNT formation. Pertinent to these findings, immunofluorescence of serum-stressed KSECs showed IQGAP1 and ARF6 expression (Figure 6A-C) within and at the base of TNTs, suggesting that CD13 again localizes these proteins to the membrane where, in this case, it facilitates TNT formation, implicating CD13 association with IQGAP1/ARF6/ in TNT formation.

In addition to inducing FAK and Src signaling, IQGAP also recruits the Rho GTPase Cdc42/Rac1-WASP-Arp2/3 complex to the plasma membrane (Rohatgi *et al*., 1999; Sugihara *et al*., 2002; Hanna *et al*., 2017), enabling the protrusion of actin polymers. Similarly, it has been shown that active Cdc42/Rac1 is critical for TNT formation in macrophages, which also express high levels of CD13 (Hanna *et al*., 2017). In agreement with these data, inhibition of Cdc42-GTP and actin polymerization, significantly reduced the number of TNTs in WT KSECs (Figure 1F).

Finally, the phosphatidylinositol species PI(4,5)P2 has been implicated in TNT formation (Kimura *et al*., 2016) and is a key regulator of actin assembly, cell membrane dynamics, focal adhesion formation and signal transduction (Mandal, 2020). ARF6 recruitment to the plasma membrane activates PIP5K (phosphatidylinositol-4-phosphate 5-kinase) to generate PI(4,5)P2 (Brown *et al*., 2001). We have shown that CD13 is important for assembling the proteins for endocytosis, and therefore hypothesized that CD13’s recruitment of ARF6 may also modulate PI(4,5)P2 generation. mAb 452 activation of CD13 in WT KSECs significantly increased PIP5K activity compared to control or serum-stressed conditions (Figure 7A), which was accompanied by a clear increase in PI(4,5)P2, indicating that activation of CD13 activates PIP5K to generate PI(4,5)P2 (Figure 7B-C). In agreement with previous work, PI(4,5)P2 localized in TNTs as visualized by transfection with PH-GFP, a plasmid encoding a labeled pleckstrin homology (PH) domain of PLCγ that specifically binds to PI(4,5)P2 (Varnai and Balla, 1998; Holz *et al*., 2000). PH-GFP/PI(4,5)P2 localization significantly correlates with that of CD13 in mAb 452-treated WT KSECs (PCC r = 0.91) in comparison to CM control (PCC r = 0.75, Figure 7D-H).

**FIGURE 7.**
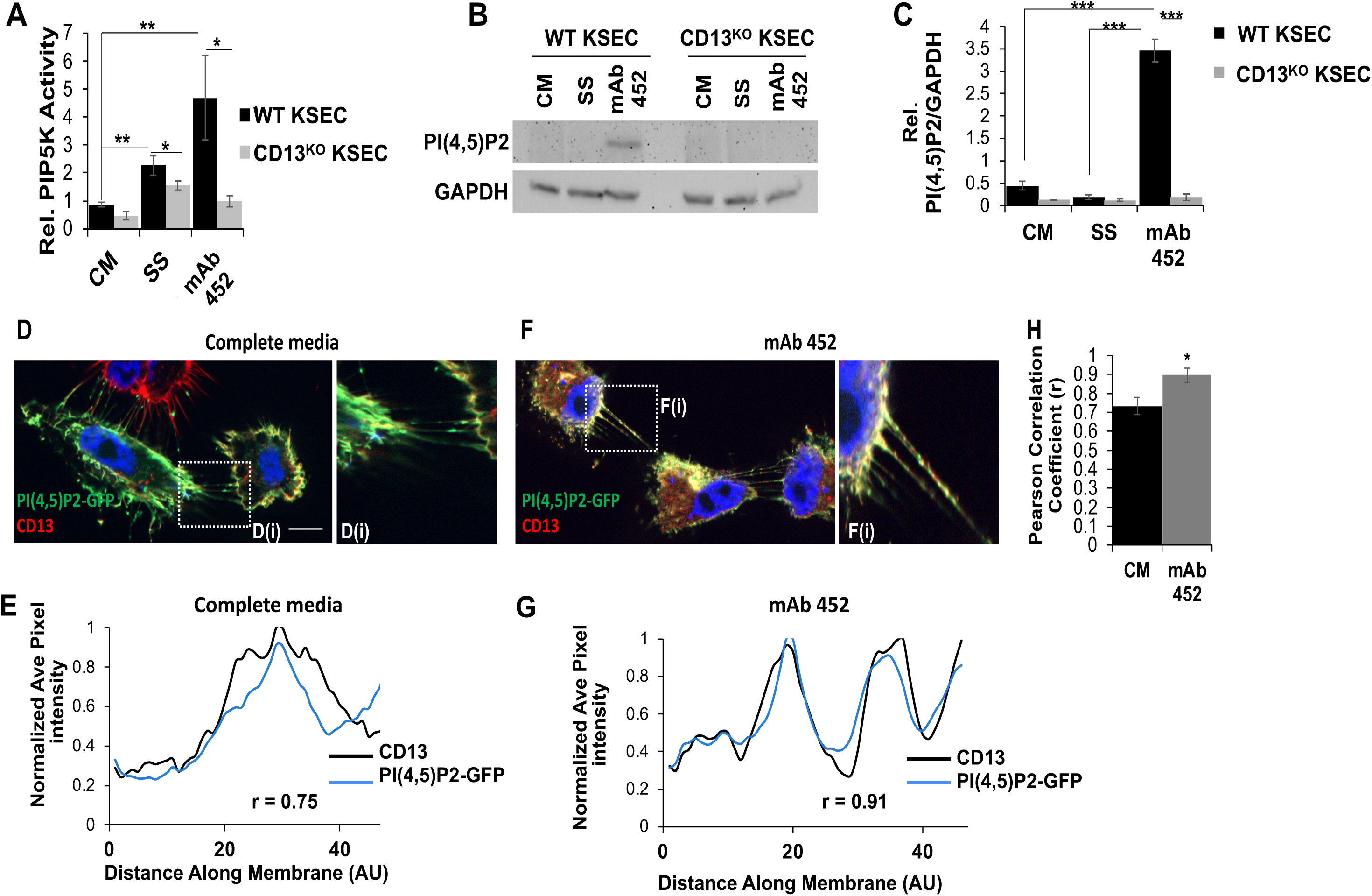
CD13 crosslinking activates PI5K and induces PI(4,5)P2 generation. **(A)** Relative PI5K activity was measured in WT and CD13^KO^ KSECs under stressed (SS) and activating (mAb 452,1.8μg/mL) conditions. **(B)** Western blot quantification of downstream PI(4,5)P2 generation under stressed and activating conditions. The undetectable PI(4,5)P2 levels in CD13^KO^ KSECs are not indicative of an absence of PI(4,5)P2 in CD13^KO^ cells, but rather suggest lower overall levels**. (D, F)** WT cells under stressed and activating conditions were stained for CD13 (red) and PI(4,5)P2-GFP (green). Enlarged image insets **(Di, Fi)** illustrate the accumulation of proteins at the base of TNTs. **(E, G)** fluorescence was quantified (CD13-black line: PI(4,5)P2-GFP blue line) using image J and Pearson’s correlation calculated **(H).** Data are represented as mean ± standard error. *p<0.05.

Together, these data describe a pathway in which activation of CD13 orchestrates FAK and Src phosphorylation, CD13-dependent recruitment of IQGAP1/ARF6 at the plasma membrane, triggering downstream Ccd42 activation and actin polymerization, PIP5K activation, ultimately resulting in the generation of PI(4,5)P2 and cytoskeletal changes required to induce TNT formation (Figure 8) (Ghosh *et al*., 2019).

**Figure 8.**
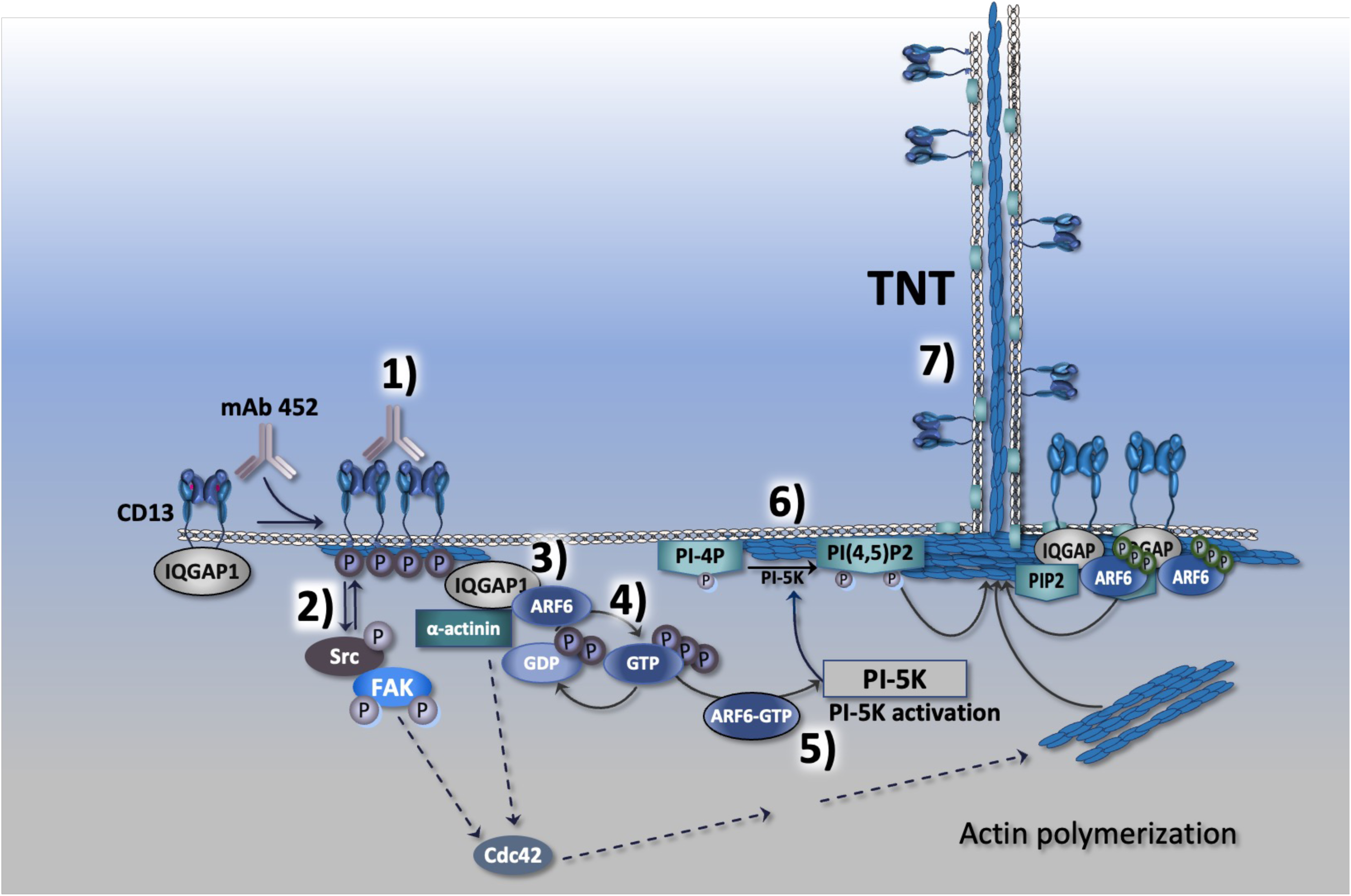
Proposed mechanism of CD13-dependent TNT formation. Crosslinking of CD13 with mAb 452 induces **1)** its phosphorylation by Src and FAK kinases, which **2)** activate Cdc42, and **3)** the assembly of a membrane-bound complex containing IQGAP and ARF6, which we propose stimulates Cdc42 and actin polymerization. In addition, by tethering IQGAP at the membrane, CD13 enables **4)** activation of ARF6 to ARF6-GTP. In turn, **5)** activated ARF6 activates PI-5K, enabling the conversion of PI-4P to PI4,5P2 **6)**, which promotes the membrane protrusion necessary for TNT formation **7).** In combination, these signal transduction pathways are responsible for CD13-activation-dependent TNT formation.

## Discussion

In the current study, we demonstrated that the cell surface molecule CD13 mediates the formation of tunneling nanotubes in a human endothelial cell line. Our studies show that CD13 acts by assembling the machinery necessary to activate signaling pathways regulating protein and lipid phosphorylation and activation, actin polymerization and membrane organization. Furthermore, we discover a novel function for CD13 in the critical membrane phosphoinositide regulatory pathway, specifically in regulating PI-5K activation of lipid phosphorylation and PIP2. Phosphoinositides impact various cellular processes via their interaction with a vast array of protein partners and other membrane phospholipids. They are themselves regulated by PI kinases and phosphatases to control the cytoskeleton, membrane dynamics and intracellular vesicular trafficking, all of which likely play a role in TNT formation and transport. Membrane PIP2 interacts with actin-binding proteins to control actin network organization (Onfelt *et al*., 2004; Wenk, 2005; Wittchen, 2009; Simons and Sampaio, 2011; Vestweber *et al*., 2013; Johnson *et al*., 2017). Pertinent to our study, PIP2 binds to α-actinin as does active CD13, likely contributing to TNT formation. Similarly, in collaboration with Cdc42, PIP2 promotes N-WASP/Arp2/3 signaling that has been implicated in actin polymerization and TNT formation (Hanna *et al*., 2017). PIP2 also regulates the focal adhesion kinase (FAK) that has been implicated in many signaling pathways controlling migration and adhesion. We have shown that the absence of CD13 reduces activation of both FAK and Src kinases and cell migration (Subramani *et al*., 2013; Ghosh *et al*., 2019) perhaps due to higher PIP2 levels upon CD13 activation. PIP2 has been shown to interact with and activate FAK in an ATP-dependent manner to mediate focal adhesion turnover and FAK/integrin signaling (Sugihara *et al*., 2002). Finally, while phosphoinositides comprise only 1% of the plasma membrane, PIP2 plays a prominent role in membrane dynamics and organization via its binding proteins. Many of these are activated by signaling induced by cell surface receptor activation (Simons and Sampaio, 2011) and contribute to membrane curvature sensing, membrane domain segregation, endocytosis and exocytosis (Pergu *et al*., 2019). We have established that CD13 is a regulator of receptor-mediated endocytosis (Ghosh *et al*., 2012; Ghosh *et al*., 2015) and integrin vesicular trafficking (Ghosh *et al*., 2019) but have not explored the contribution of phosphoinositides to these CD13 functions. Further investigation into how these multifunctional proteins interrelate in these cellular processes will clearly uncover novel common mechanisms regulating TNT formation.

In contrast to its developmentally regulated expression in myeloid cells during hematopoiesis, CD13 is not expressed in resting endothelial cells but is induced and activated in response to growth factors and signals elicited under pro-angiogenic and - inflammatory conditions (Bhagwat *et al*., 2001; Bhagwat *et al*., 2003; Petrovic *et al*., 2003). Interestingly, TNTs have been implicated in both inflammation (Jiang *et al*., 2016; Weng *et al*., 2016; Ariazi *et al*., 2017; Nawaz and Fatima, 2017; Weidle *et al*., 2018; Yao *et al*., 2018; Kuret *et al*., 2019; Mittal *et al*., 2019a) and angiogenesis (Figeac *et al*., 2014; Climent *et al*., 2015; Patheja and Sahu, 2017; Errede *et al*., 2018; Lou *et al*., 2018; Liu *et al*., 2019), indicating that CD13 may be upregulated specifically to participate in these processes, possibly by promoting TNT formation and function. We have also determined that during inflammation, activation of CD13 on endothelial cells increases monocyte adhesion and transmigration (Mina-Osorio *et al*., 2008; Subramani *et al*., 2013; Ghosh *et al*., 2014; Rahman *et al*., 2014a; Rahman *et al*., 2014b), which may relate to “transmigratory cups” or “endothelial adhesive platforms” which are actin-based, TNT-like docking protrusions that form at the site of inflammatory leukocyte extravasation to mediate the adhesion phase of immune infiltration (Carman *et al*., 2003; Carman and Springer, 2004; Riethmuller *et al*., 2008; Wittchen, 2009; Vestweber *et al*., 2013; Franz *et al*., 2016; Johnson *et al*., 2017; Genna *et al*., 2023). In support of this notion, we have found that endothelial CD13 is highly enriched in similar protrusions formed between monocytes and endothelial cells in vitro (Mina-Osorio *et al*., 2008). Similarly, the hypoxic environment of tumors potently induces endothelial expression of angiogenic growth factors and CD13 (Bhagwat *et al*., 2001) and TNT formation (Lou *et al*., 2018), again suggesting that promoting TNT formation may be a component of CD13’s overall regulation of endothelial invasion. Similar to CD13, the CSPG4 protein (also known as NG2) is a transmembrane angiogenic regulator (Chekenya *et al*., 2002) that is also upregulated in response to hypoxia and inflammatory signals (Ampofo *et al*., 2017) and participates in the activation of various signaling pathway components such as FAK, Erk 1, 2 and Rho family GTPases by virtue of its cytoplasmic tail (Price *et al*., 2011). Interestingly, overexpression of NG2 in cancer cell lines stimulates TNT formation (Lou *et al*., 2018), consistent with inducible angiogenic regulators as potential mediators of TNT function. It will be interesting to investigate CD13 dependent TNT formation in heterotypic co-cultures of myeloid and endothelial cells or tumor cells (Genna *et al*., 2023).

Accumulating evidence demonstrates that not surprisingly, TNT formation and cargo transfer requires contributions from a large number of proteins of various functions including GTPases [RalA (Hase *et al*., 2009; Schiller *et al*., 2013)], vesicular tethering, transport and recycling proteins [Rabs 8 and 11 (Zhu *et al*., 2018), M-Sec (Hase *et al*., 2009)), cytoskeletal regulators [RASSF1A (Dubois *et al*., 2018)], actomyosin-motor proteins [myo10 (Gousset *et al*., 2013; Reichert *et al*., 2016)] and ER chaperones [ERp29 (Pergu *et al*., 2019)] among others. Logically, these components must be carefully assembled into multimolecular complexes by scaffolding proteins and localized to specific areas of the cell to enable these processes. One such multimolecular complex has been identified to coordinate TNT formation, where overexpression of the membrane-bound LST1 (lymphocyte specific transmembrane protein) in HeLa cells induces TNT formation by recruiting a complex containing the small GTPase RalA, its GEF RalAGPS2, the actin-binding scaffold protein filamin and the exocyst-homologous protein M-Sec to the plasma membrane (Schiller *et al*., 2013). We have recently described a similar role for CD13 in the recruitment and tethering of a complex containing analogous components; the GTPase ARF6 and its GEF EFA6, the scaffold protein IQGAP1 and actin-binding α-actinin at the plasma membrane to regulate β1-integrin endosomal recycling (Ghosh *et al*., 2019). Interestingly, these complexes may be functionally linked: integrins have been shown to activate RalA (a component of the LST1 complex) that in turn stimulates ARF6 (a component of the CD13 complex) to drive exocyst-dependent delivery of membrane microdomains to the plasma membrane (Pawar *et al*., 2016), perhaps to supply membrane to the growing TNT. In agreement with this mechanism, we find that in CD13^KO^ KSECs (which are impaired in TNT formation) ARF6 is in its inactive GDP form and β1-integrin accumulates in cytoplasmic vesicles rather than recycling to the membrane, perhaps interfering with integrin-Ral-ARF6 activation and subsequent membrane expansion. Furthermore, IQGAP1, β1-integrin, FAK and Src proteins are found along the length of the TNT (not shown), consistent with a functional role for this CD13-tethered recycling complex in TNT formation, which is under current investigation.

In addition to facilitating TNT formation, we have recently reported that CD13 also negatively regulates cell-cell fusion, in part by limiting the formation of fusion-promoting, actin-based protrusions located at the base of the cell (Ghosh *et al*., 2024). Specifically, we found a significant increase in micro protrusion-like structures (visible under electron microscopy) attached to the substratum in CD13^KO^ macrophages under fusion-promoting conditions. These structures contributed to exaggerated macrophage fusion to form the multinuclear giant cells (MGCs) that characterize the inflammatory reaction induced by foreign implants (Carnicer-Lombarte *et al*.). While this observation may seem at odds with our current study describing CD13 as a stimulator of TNT formation, we find that KSECs lacking CD13 also show increased numbers of short protrusions at the base of the cell (data not shown). The length of these structures and their location at the base of the cell suggests that they are similar to filopodia and not TNTs, although further characterization is needed. It is possible that under cellular stress, CD13 is a molecular switch that promotes the formation of TNTs for exchange of cargo to rescue of distant cells, over filopodia. Further investigation into CD13 and its effects on actin dynamics may help us to distinguish among the molecules and machinery that regulate filopodia, fusion-promoting protrusions, and TNTs.

Finally, while this work identifies a unique CD13-dependent molecular complex controlling TNT formation, it is also novel in that it defines CD13 activation as one of very few molecular triggers that induce the formation of functional TNTs. Combined with similar novel observations, our findings will facilitate the dissection of the pathways, molecules and mechanisms that underlie TNT formation and contribute to a standard to enable classification of these and other actin-based structures (Belian *et al*., 2023).

## Methods

### Cell Culture

Kaposi Sarcoma (KS1767) endothelial cells were cultured in DMEM (Gibco 11965-092) supplemented with 1% Penicillin-Streptomycin (P/S) (Gibco 15140122, 100U/mL) and 10% HI-FBS (Gibco 16250078). Cells were detached using 5mL 0.25% Trypsin-EDTA (Gibco 25300054), collected into 10mL fresh serum-containing DMEM, spun at 1200rpm for 5mins and counted using trypan blue and a Countess II. For the continuation of cells in culture, 2×10^6^ cells were seeded per T175 flask with 25mL DMEM supplemented with P/S and HI-FBS. For experiments, cell seeding density varied depending on the application from 1×10^5^-3×10^5^ cells/coverslip. Cells seeded onto coverslips were resuspended in DMEM supplemented with P/S and HI-FBS. Cells then settled overnight, or for 2-3hrs, depending on the application, prior to treatments.

Human umbilical vein endothelial cells (HUVECs) were cultured in EBM-2 (Lonza CC-3156) media supplemented with EGM-2 SingleQuots Supplements (Lonza CC-4176) until 80% confluent. Cells were detached using 5mL 0.05% Trypsin-EDTA (Gibco 25300054), collected into 10mL fresh serum-containing media, spun at 1200rpm for 5mins and counted using trypan blue and a Countess II. For the continuation of cells in culture, 2×10^6^ cells were seeded per T175 flask with 25mL EBM-2 media. For experiments, cell seeding density varied depending on the application from 1×10^5^-2×10^5^ cells/coverslip. Cells settled overnight onto coverslips prior to treatments.

### Antibody and Ligand Treatment of Cells

KSECs and HUVECs were seeded at a density of 7.5×10^4^ cells/coverslip. Cells attached to coverslips overnight and washed two times with PBS followed by the addition of any treatments for 1hr. (see immunofluorescence section for more details.)

| Treatment | Working Concentration | Catalogue # |
| --- | --- | --- |
| CD13 mAb | 1.8µg/mL | Isolated by Shapiro Lab |
| Bradykinin | 200nM | Sigma B3259 |
| WM4.7 | 0.1µg/mL | Sigma C8589 |
| CD31 | 1:1000 (1µg/mL) | Invitrogen MA3100 |
| Bestatin | 100µg/mL | Sigma B8385 |
| Cytochalasin B | 250nM | Cayman Chemical 11328 |
| Cytochalasin D | 250nM | ThermoFisher PHZ1063 |
| FAK14 | 20µM | Sigma SML0837 |
| PP2 | 10µM | Selleck Chem S7008 |
| ML141 | 20µM | Tocris 4266 |
| Nocodazole | 150nM | Selleck Chem S2775 |

### Immunofluorescence and Labeling

| Antibodies/stains | Dilution/concentration, catalogue # |
| --- | --- |
| CD13 mAb | 1:100 (1.8µg/ml), isolated |
| Phalloidin-TRITC | 1:100 (1µg/mL), Sigma #P1951 |
| Phalloidin-488 | 1:1000 (0.3U/mL), ThermoFisher #A12379 |
| Alexa Fluor secondary antibodies (Life Technologies) | 1:1200 (1.6µg/mL)<br>488: #A11029, #R37114<br>594: #A11037, #A21203<br>640: #A21235 |
| DAPI | 1:500, Invitrogen #D1306 |

Cells were washed two times with PBS, fixed for 20mins in 4% PFA at room temperature and permeabilized in 0.01% Triton-X for 5mins at room temperature. Coverslips were then blocked for 1hr at room temperature in PBS/5% w/v BSA supplemented with 5% serum from the host of the secondary antibody. Primary antibodies were added (see list above) in the same PBS/5%BSA/5%host serum and coverslips were incubated overnight at 4 °C. The cells were then washed twice with PBS and incubated for 1 hour at room temperature with Alexa Fluor secondary antibodies (see above) and DAPI. The coverslips were then washed three times with PBS and mounted upside down on glass slides with Prolong Gold Antifade mounting media (1 drop, Molecular Probes P10144). The mounting media was left to cure overnight at room temperature and protected from light.

### Microscopy Imaging

Slides were imaged using either an inverted fluorescent Zeiss LSM880 microscope or a confocal Zeiss Axio Observer.Z1 with a 63X oil objective. Images and Z-stacks were then taken using the ZEN 2 software by Zeiss and scale bars were added.

#### Image Analysis

TNT index was calculated as an average number of TNTs/eligible cell. Cells were considered eligible for analysis if at least 90% of the cell body was within the FOV, there was sufficient room surrounding the cell to form a TNT (e.g a cell of interest could not be counted if it was touching other cells on all sides). Total TNTs from eligible cells were then counted per FOV and normalized to the number of total eligible cells to get the TNT index.

### Measurement and correlation of protein co-localization by immuno-fluorescence

In graph overlay, the x-axes represent the relative measurement position from the arbitrary initial point #1 along the entire cell membrane. The values on the y-axes were obtained by dividing each intensity measurement by the overall maximum pixel intensity in that cell to obtain a normalized relative pixel intensity at each point along the cell membrane. Similar curves represent similarities in expression distribution as verified by Pearson Correlation Coefficient (PCC) calculation. PCC analysis was performed to determine the correlation of pixel intensity measurements for CD13 and phalloidin.

### Ca+2 flux measurement

Changes of intracellular Ca^2+^ was measured using ratio Ca^2+^ imaging as described previously(Zong *et al*., 2022). In brief, Fura-2 AM (Thermal Fisher Scientific, F1221) was dissolved in DMSO to make a stock concentration at 1 mM. Pre-warmed Neurobasal® Medium (Thermal Fisher Scientific, 21103-049) was used to dilute Fura-2 AM to a working concentration at 2.5 µM, and 0.02% Pluronic™ F-127 (Thermal Fisher Scientific, P3000MP) was added to facilitate loading of Fura-2 AM. Cells plated on 25 mm glass coverslips were washed using pre-warmed PBS for 3 times, and then incubated with 2 ml of Fura-2 AM working solution for 30∼45 min at 37 °C. Non-incorporated dye was washed away using HEPES-buffered Saline Solution (HBSS) containing (in mM): 20 HEPES, 10 glucose, 1.2 MgCl_2_, 1.2 KH_2_PO_4_, 4.7 KCl, 140 NaCl, 1.3 Ca^2+^ (pH 7.4). Patch electrodes were pulled from borosilicate glass and fire-polished to a resistance of ∼3 MΩ when filled with HBSS solutions.

### Statistical Analysis

Statistical analysis was performed using unpaired, two-tailed Student’s t test and results are representative of mean ± SEM as indicated. Differences at p≤ 0.05 were considered significant.

---

**FIGURE S1:**
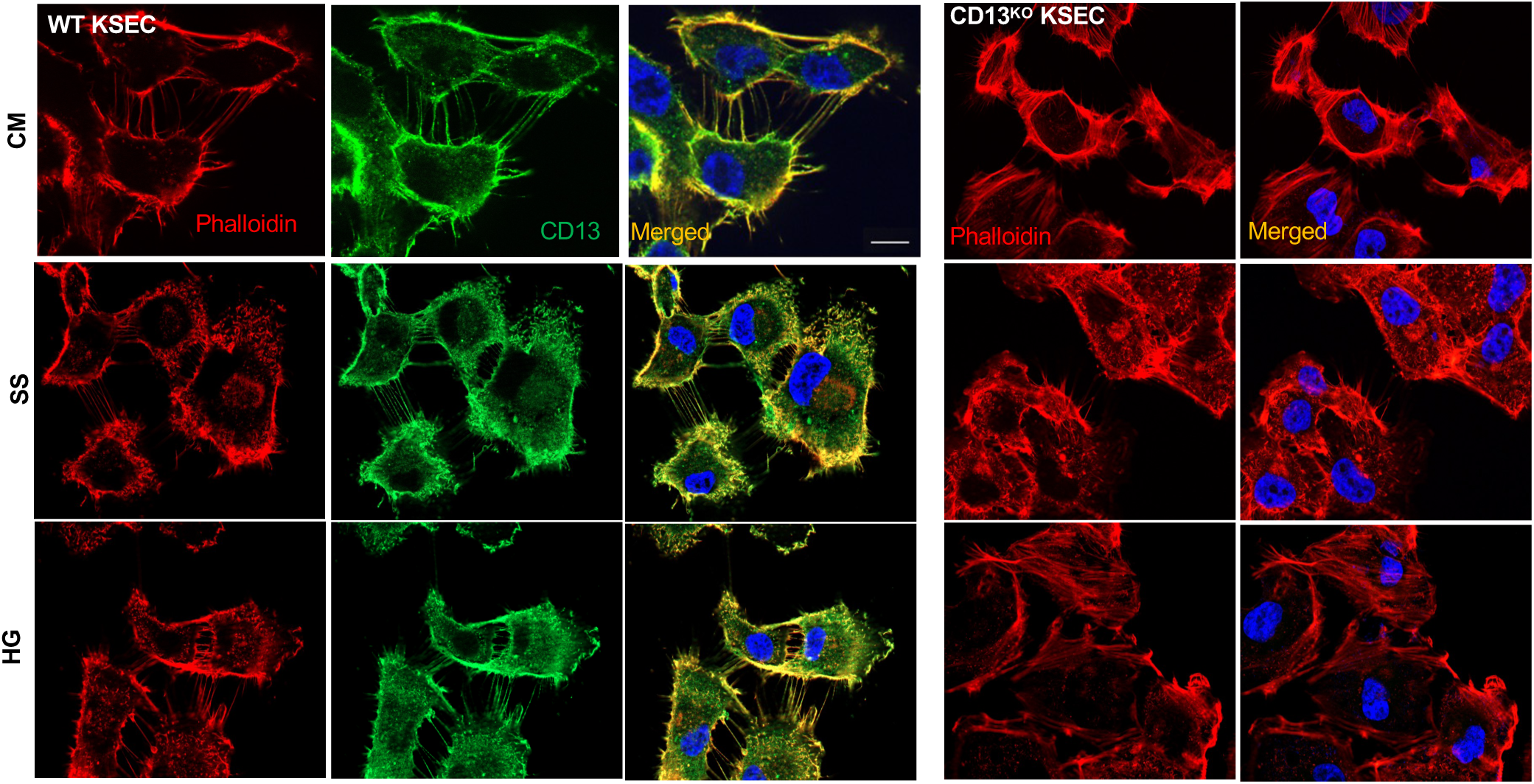
Split-channel comparison of WT and CD13KO KSECs forming TNTs. Phalloidin (red) and CD13 (green) channels separated across rows followed by merged images with DAPI (blue) for WT and CD13KO KSECs. Rows represent culture conditions with complete media (CM), serum starvation (SS) or high glucose (HG). Scale bar: 10μm.

**FIGURE S2:**
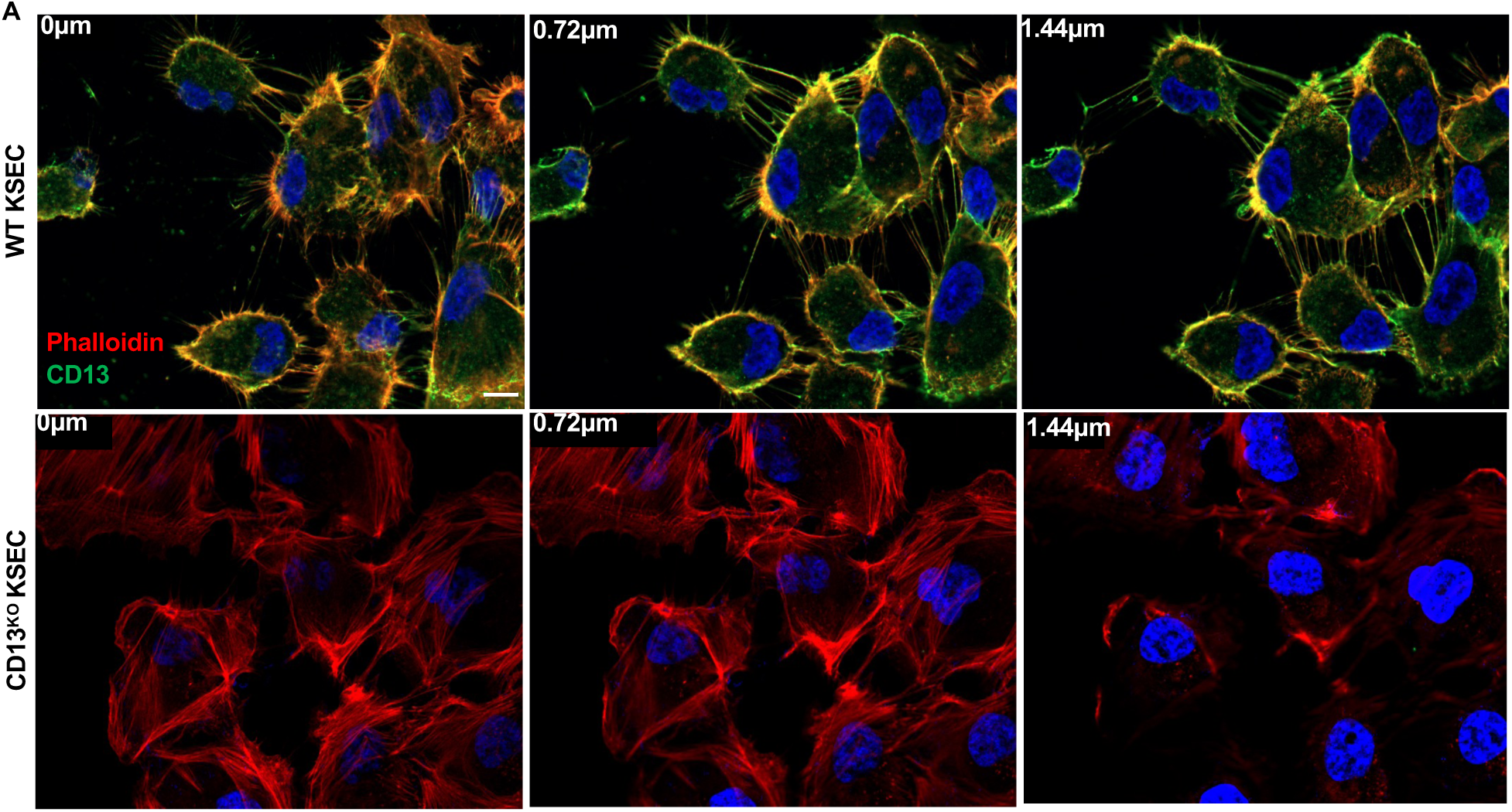
Protrusions in WT cells form further from the substrate than in CD13KO cells. (A) representative z-stack images of WT and CD13KO cells with protrusions spanning between 0.48-1.44μm above the substrate in WT KSECs. Scale bar: 10μm.

**FIGURE S3:**
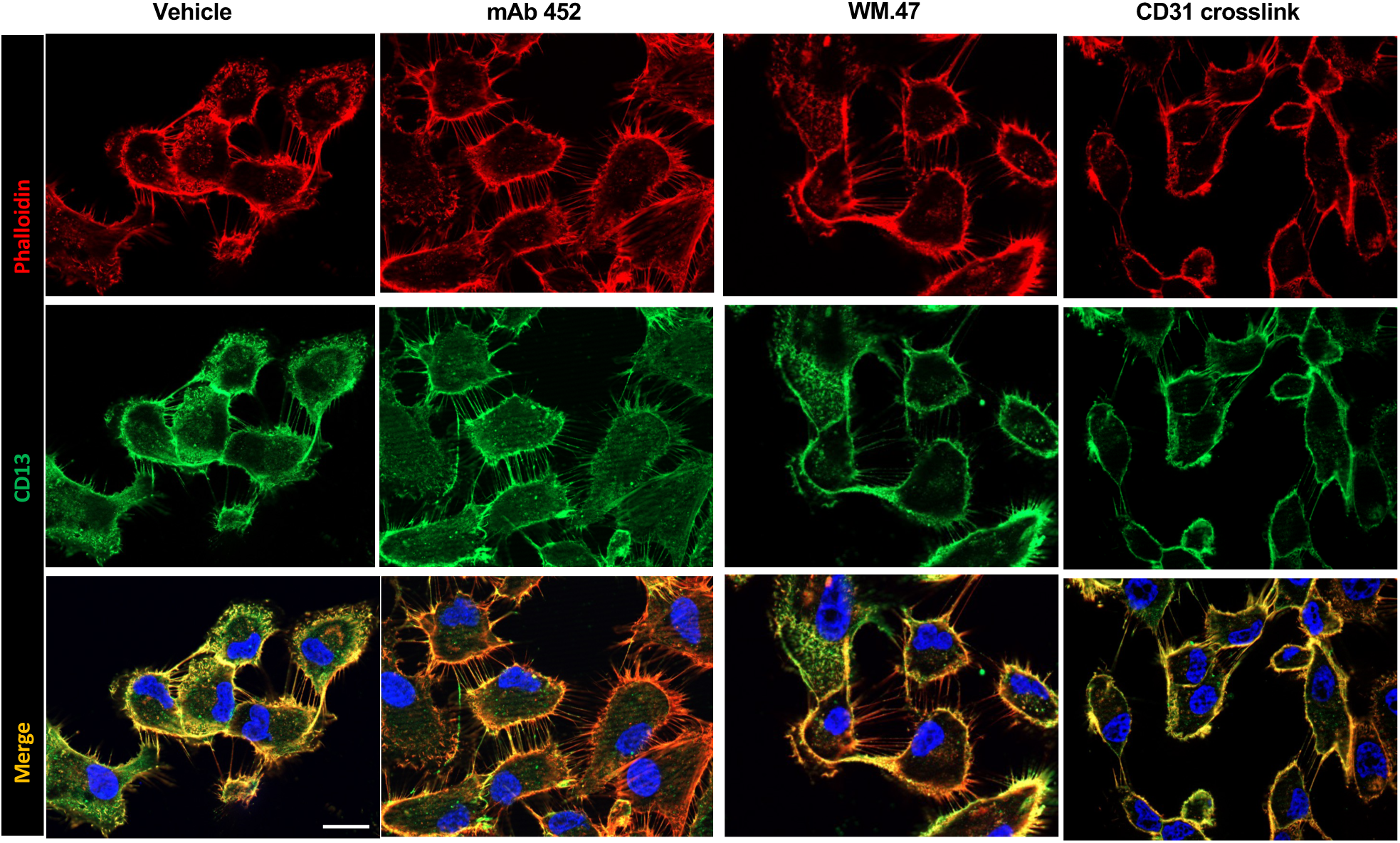
Split-channel comparison of CD13 and phalloidin staining in WT KSECs treated with crosslinking or non-crosslinking antibodies. Phalloidin (red) and CD13 (green) channels separated across rows followed by merged images with DAPI (blue) for WT KSECs. Rows represent culture conditions with vehicle control (Vehicle; complete media), mAb 452 (1.8μg/mL), CD13 mAb WM4.7 (1:100), or CD31 crosslink (1μg/mL). Scale bar: 10μm.

**FIGURE S4:**
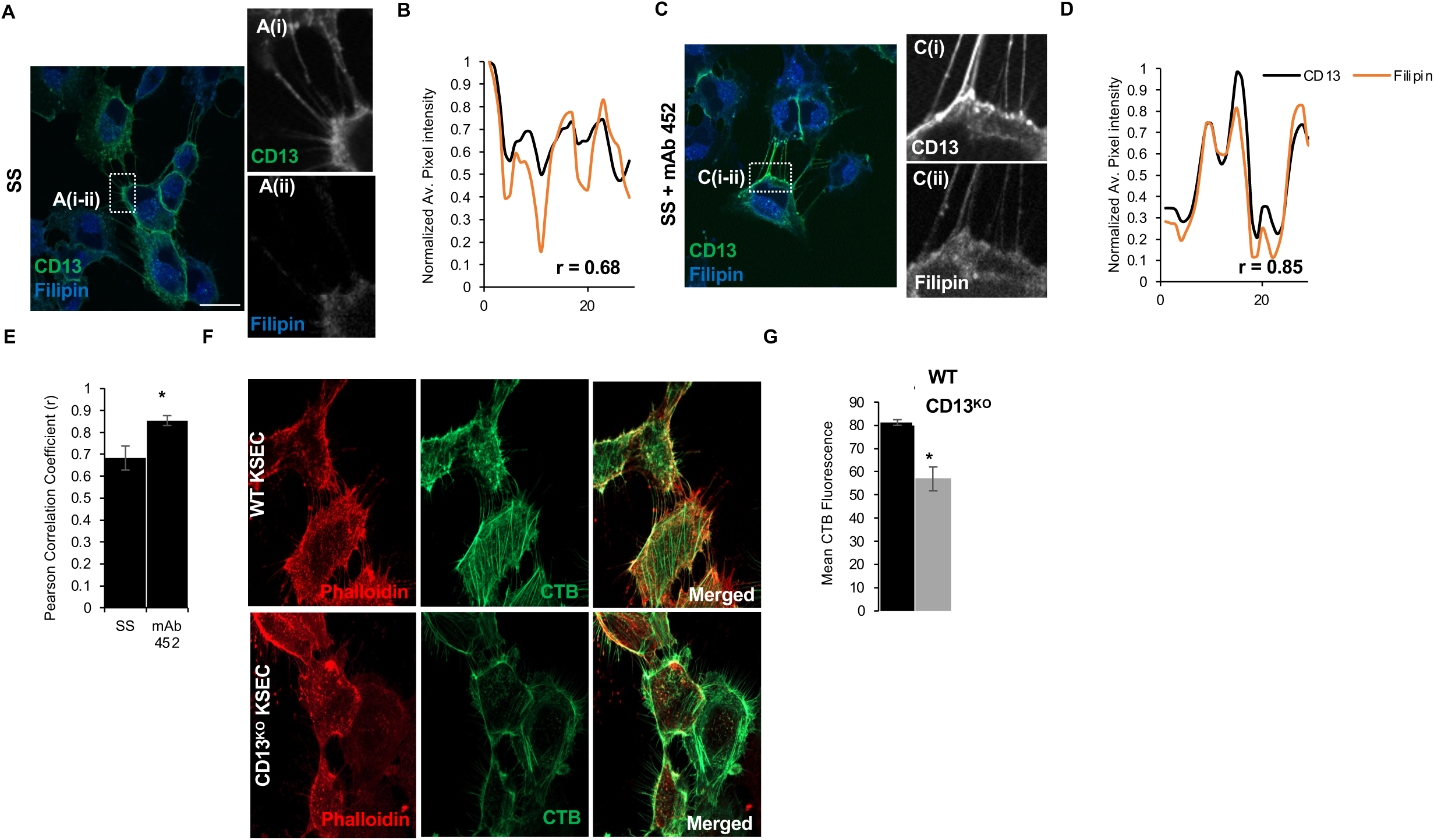
CD13 may localize membrane lipids at the base of TNTs. (A-D) Representative images and overlaid membrane line scans for localization of CD13 (green) and filipin (blue) at the base of TNTs in WT KSECs in serum-starved (A, B; SS) or mAb 452 (C, D; 1.8μg/mL) treatment conditions. (E) Comparison of the average Pearson correlation coefficient in cells under SS or mAb 452 conditions. (F) Representative images of CTB staining in WT versus CD13KO KSECs. (G) quantification of mean CTB fluorescence on WT versus CD13KO KSECs. Scale bar: 10μm. Data are represented as mean ± standard error. * = p<0.05.

**FIGURE S5:**
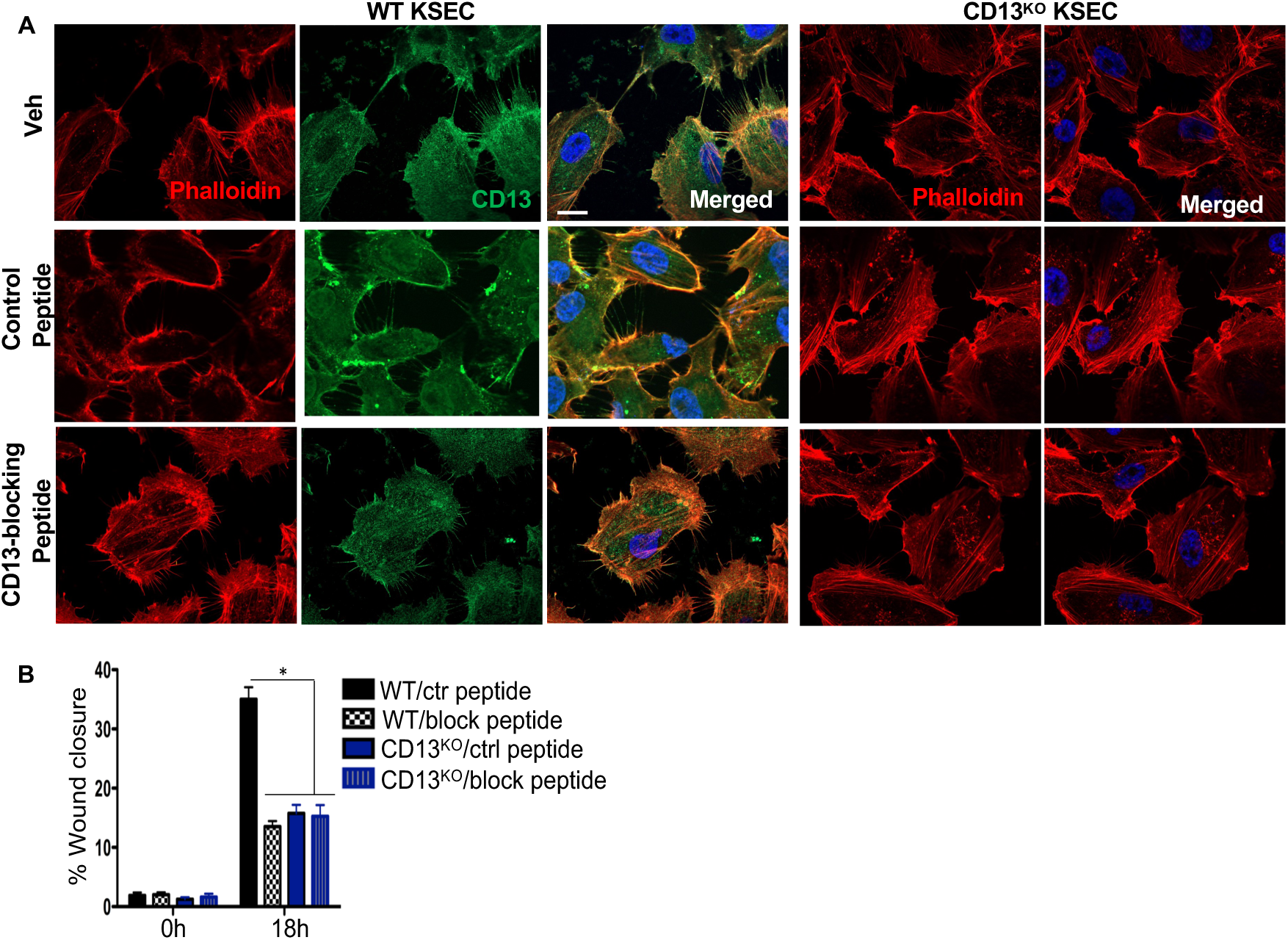
CD13-blocking peptide effectively mimics KO in wound-closure assay. (A) Images of separated channels of WT and CD13KO KSECs, phalloidin (red), CD13 (green) merged with DAPI (blue) of vehicle control, control (1μg/mL) or CD13-blocking (1μg/mL) peptide loaded cells. (B) CD13 has been shown to regulate cell migration during wound healing. Blocking CD13 with this peptide significantly inhibited wound closure, like in CD13KO KSECs. Data are represented as mean ± standard error. *p<0.05. Scale bar: 10μm.

**FIGURE S6:**
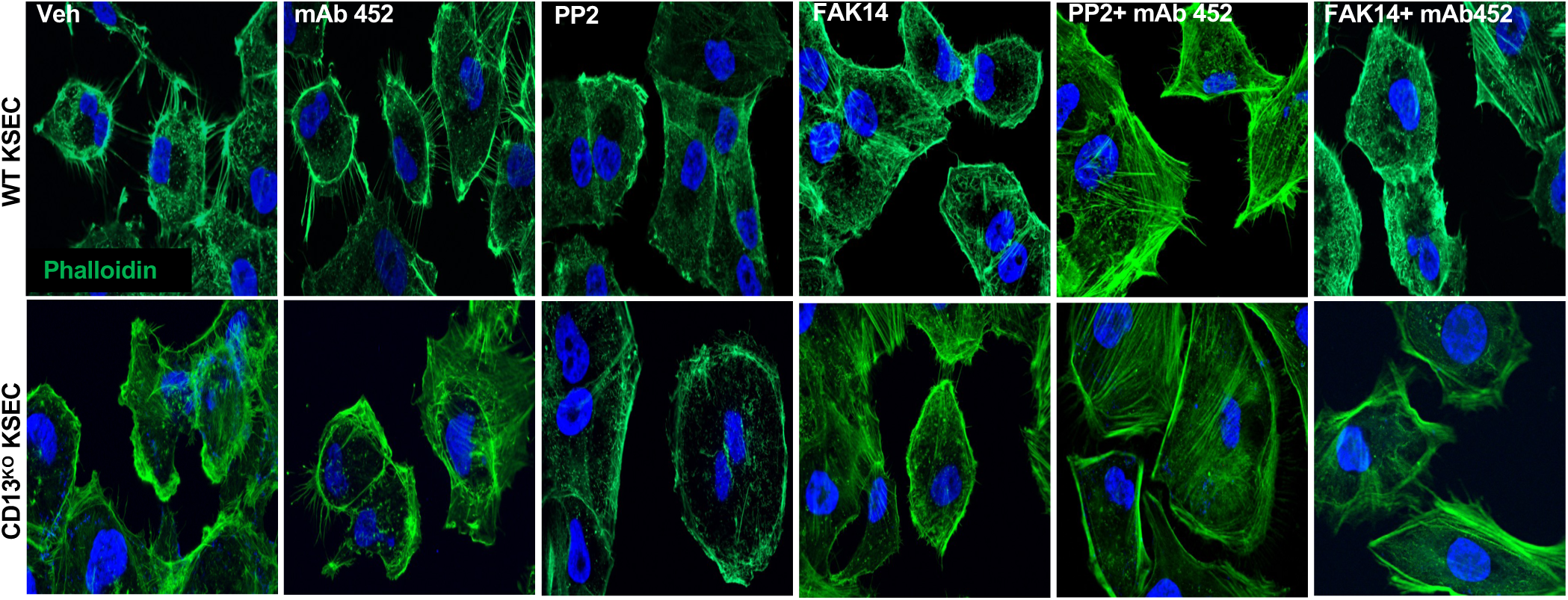
Inhibition of FAK/Src signaling abrogates TNT formation. WT KSECs were treated with a FAK inhibitor 14 (FAK14; 20μM) or Src inhibitor (PP2; 10μM) with or without mAb 452(1.8μg/mL), for 1hr prior to staining with phalloidin (green) and DAPI (blue).

